# Patchwork: alignment-based retrieval and concatenation of phylogenetic markers from genomic data

**DOI:** 10.1101/2022.07.03.498606

**Authors:** Felix Thalén, Clara G. Köhne, Christoph Bleidorn

## Abstract

**Motivation:** Increased output from the latest short-read sequencers makes low-coverage whole-genome sequencing (LC-WGS) an increasingly affordable approach to large-scale phylogenetics. Despite offering several advantages over prevailing sequencing strategies, few tools exist to work with this data type within a phylogenomic context. Due to the fragmented nature of LC-WGS genomes, their use have mostly been restricted to easy-to-assemble, high-copy-number regions such as organelle genomes and or ribosomal genes.

**Results:** We here present a new method for mining phylogenetic markers directly from an assembled genome. Homologous regions are obtained via an alignment search, followed by a “hit-stitching” phase, in which adjacent or overlapping regions are concatenated together. Finally, a novel sliding window technique is used to trim non-coding regions from the alignments. We demonstrate the utility of Patchwork by recovering near-universal single-copy orthologs (USCOs) in the annelid *Dimorphilus gyrociliatus.*

**Availability:** Patchwork is available from Github under the GNU General Public license version 3.

**Supplementary information:** Supplementary data are available at github.com/Animal-Evolution-and-Biodiversity/benchmarking-patchwork.

## 1 Introduction

Advancements in high-throughput sequencing techniques have revolutionized the field of phylogenetics and ultimately our understanding of the tree of life (Lemmon et al, 2012). The availability of genomic and or transcriptomic data for basically all desired taxa and for a reasonable price has transformed the field to phylogenomics— genome-scale phylogenetic systematic analyses (McCormack et al, 2013). Some challenges remain, however, as many studies still show incongruent results, lack of branch-support, or resolution (Philippe et al, 2017). Even though complete genomes are available for more and more eukaryotes, most large-scale phylogenomic studies to date were conducted using either transcriptome sequencing (e.g., Weigert et al, 2014; Andrade et al, 2015) or a genome subsampling methods such as targeted-enrichment (e.g., Andermann et al, 2020; Sann et al, 2018; Call et al, 2021) which focuses on a set of pre-selected loci.

### 1.1 Reduced representation vs. WGS strategies

Transcriptome sequencing offers a way to sequence only the expressed portion of a genome without prior sequence knowledge. Unfortunately, this approach requires freshly collected material or material stored in a specific manner (e.g., deeply shock-frozen, RNAlater; Cronn et al, 2012). Furthermore, smaller specimen may need to be pooled together to attain sufficient amounts of mRNA and such practice risks mixing up individuals with undetected genetic variation (Allen et al, 2017). Unfortunately, a large amount of collected specimen only exist in natural history museum collections and most of these are ethanol-preserved and thus not usable for transcriptomic studies (Call et al, 2021). This is undesirable as taxon sampling is considered one of the most important factors for accurate phylogenetic tree reconstruction (Heath et al, 2008).

Target-enrichment approaches, on the other hand, require prior knowledge of target sequences (e.g., from well-annotated genomes) for the construction of oligonucleotide probes. Moreover, the number of enriched targets are limited by the amount of oligonucleotides included in the enrichment kit of choice and the efficiency of such approaches decreases as the distance bait-to-target distance increases (Bragg et al, 2016). Another downside is that the data produced have few applications outside of phylogenomics and if one wants to add a taxon to the study, they need to use the exact same markers as previous studies (Allen et al, 2017).

A viable alternative is low-coverage whole genome sequencing (LC-WGS; also known as “shallow genome sequencing”, or “genome skimming”) using short-read techniques such as Illumina sequencing. Relying solely on this technique has been shown to be inadequate for the reconstruction of highly contiguous reference-quality genomes (Rhie et al, 2021). However, due to the introduction of newer sequencing platforms (e.g., Illumina’s NovaSeq sequencing platform) WGS became relatively cheap (Schwarz et al, 2021) and even highly fragmented DNA can be used as input. Consequently, LC-WGS can be used to generate data from various sources of targeted organisms to retrieve marker loci on a genome scale. While this so called “genome-skimming” approach has frequently been used to reconstruct organellar genomes (e.g., Jin et al, 2020; Richter et al, 2015), it is currently underutilized in the field of phylogenomics.

Short-read assemblies of eukaryotic genomes tend to be highly discontinuous and automated annotation of such large, fragmented genomes remains difficult (Salzberg, 2019). Eukaryotic genomes are characterized by the presence of “genes in pieces”, where introns interrupt coding sequences and exons (Rogozin et al, 2005). Depending on the coverage, short-read draft genomes are characterized by low N50s in the range of few kbp (if at all) (Salzberg et al, 2012) and consequently, exons of a single gene usually end up on several contigs in fragmented genomes.

### 1.2 Marker discovery techniques

The disuse of genome skimming in large-scale phylogenetics could potentially be ascribed to the lack of suitable data analysis methods (Zhang et al, 2019; Philippe et al, 2011). Existing software tools for working with WGS data in a phylogenomic context, such as aTRAM (Allen et al, 2018), ALiBaSeq (Knyshov et al, 2021), and GeMoMa (Keilwagen et al, 2016, 2018), are either difficult-to-use and or written in an interpreted language (e.g., Perl or Python) which does not allow the program to scale well with the large biological datasets that are commonplace today (Knyshov et al, 2021).

To address the limitations typically associated with LC-WGS, we present Patchwork, an alignment-based tool for mining phylogenetic markers, directly from WGS data. Patchwork utilizes the sequence aligner DIAMOND (Buchfink et al, 2021), and is written in the programming language Julia (Bezanson et al, 2017), to achieve the best possible speed, thus allowing Patchwork to scale well with today’s genome-scale datasets. In addition, our implementation focuses on ease-of-use; while pre-existing methods may require the user to perform the sequence alignments separately, our program handles each step in the analysis—from start to finish.

## 2 Implementation

Patchwork is a reference-and alignment-based method for mining phylogenetic markers from WGS data. One or more reference protein sequences guide the “stitching” process, where the best-scoring, translated query nucleotide sequences are merged into continuous stretches of amino acid sequences. Merged sequences go through a masking step, where unaligned residues, ambiguous amino acid characters, and stop codons are removed from query sequences. Finally, Patchwork implements a sliding-window based alignment trimming step to rid the resulting sequences from poorly aligned residues, e.g., due to putative non-coding regions. The aim of Patchwork is to capture multi-exon or fragmented genes, scattered across different contigs in an assembled genome. Moreover, this method allows sequences of different data types (i.e., genomic and transcriptomic data) to be combined into a single dataset.

The core of Patchwork is implemented in the relatively new scientific programming language Julia. Julia strives to be as performant as possible, while still retaining a high level of productivity. Existing libraries such as BioAlignments.jl (https://github.com/BioJulia/BioAlignments.jl) and BioSequences.jl (https://github.com/BioJulia/BioSequences.jl) further sped up the development itself. Patchwork is obtainable from GitHub (https://github.com/fethalen/Patchwork), it is released under the GPLv3 license and targets both Linux and macOS. To make the installation of Patchwork easier, we also provide a Docker image that contain Julia, Patchwork, and DIAMOND, one of the external dependency.

## 3 Algorithmic Overview of Patchwork

Patchwork’s workflow can be divided into five different steps: (i) pooling of reference sequences, database construction, and initial alignment, (ii) hit stitching, (iii) alignment masking, (iv) alignment trimming, and (v) final alignment, filtering, statistical reports, and plots.

### 3.1 Initial alignment and database construction

First, all reference protein sequences—regardless of whether they are spread across multiple FASTA files or not—are pooled together into a single FASTA file, from which a DIAMOND database is created. There is also the option to use an existing DIAMOND-formatted database or a BLAST output file in a tabular format by using the `--database` or `--tabular` option respectively. These files are both provided in the output of Patchwork and can thus be re-utilized when trying out different parameters. In either case, DIAMOND’s BLASTX algorithm is used to align translated nucleotide sequences to one or more reference protein sequences.

Like DIAMOND, Patchwork, by default, scores alignments using the substitution matrix BLOSUM62 (Henikoff and Henikoff, 1996), a gap open penalty of 11, and an extension penalty of 1. Other—built-in or custom—substitution matrices may be used in place of the default option. User-chosen gap open-and gap extension penalties may also be chosen, as long as they are within the limits set by the substitution matrix of choice. DIAMOND-specific options may be set using the `--diamond-flags` option. For the user’s convenience, flags such as `--evalue` for changing the maximum expected value. Lastly, DIAMOND sensitivity modes (e.g., `--very-sensitive` and `--ultra-sensitive`), Buchfink et al, 2021, all have easy-to-access options as well.

Since the alignment search is likely to result in more than one hit, certain measures are taken to ensure that none of these hits are overlapping: They are, “hit stitching” (also known as contig-or exon stitching; i.e., merging of overlapping regions), removal of unaligned residues, and concatenation of non-overlapping regions. A graphical overview of these can also be seen in figure 1.

**Fig. 1.**
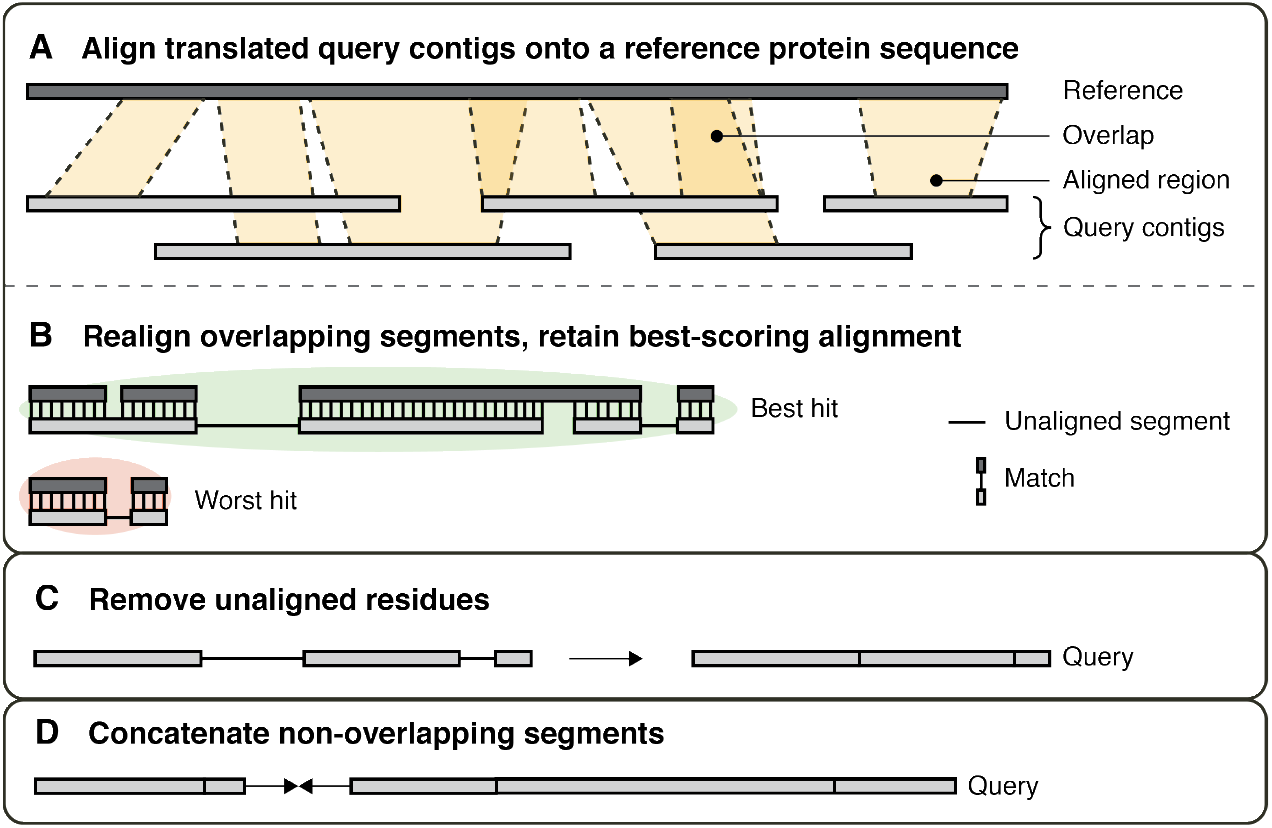
Graphical overview of the Patchwork algorithm. First, (A) query sequences are aligned to the provided reference sequence. These alignments may or may not be overlapping. (B) Overlapping alignments are realigned but only in the area in which they overlap. The best-scoring alignment is retained while all others are discarded. (C) Non-aligned residues are then removed and (D) the remaining regions are concatenated into a single, continuous sequence.

### 3.2 Hit stitching

The merging algorithm works as follows: first, all regions are sorted by their first *and* last position at which they align to the reference sequence. The first region is added to the stack and then for each pair of regions, check if the two regions are overlapping. If they are not overlapping, add the rightmost region to the stack and continue. If they are overlapping, however, realign the overlapping region to identify the best-scoring sequence at that particular interval. Then, based on the realignment score, slice the sequences such that the best-scoring sequence stays at the overlapping region and so that non-overlapping, flanking regions, if existing, are retained as well.

Different aligned regions from the same contig are allowed to be stitched together. While “hit stitching” may result in the creation of chimeric sequences (i.e., two or more biological sequences incorrectly joined together), this procedure has the potential to increase coverage and to (correctly) join two or more regions that are located on separate contigs due to an erroneous assembly, a sequencing error, and or a multi-locus gene.

### 3.3 Alignment masking

At this step, unaligned residues, ambiguous amino acid characters, and stop codons (also known as “termination codons”) are all removed from the resulting query sequence. Query sequences may contain residues which do not align to any particular region of the subject sequence. Such regions may, e.g., be non-coding regions or simply insertions. In either case, unaligned residues are removed on the basis that inserts are less likely to constitute phylogenetically informative sites and risks introducing untranslated regions and therefore biasing the downstream analysis. Similarly, ambiguous amino acids are most likely non-informative and stop codons are a clear indicator that non-coding characters have been included in the alignment. Although such regions are likely to be removed in the subsequent step (see section 3.4), the user may choose to keep stop codons and or ambiguous amino acid characters by providing the flags `--retain-stops` and or `--retain-ambiguous`.

### 3.4 Sliding window-based alignment trimming

One side effect of aligning translated nucleotide sequences to amino acid sequences is that one might recover noncoding portions of DNA, provided that the following two conditions are fulfilled: (i) the noncoding DNA is located in between two or more coding portions and (ii) there is a sequence region in the reference sequence that the noncoding region can align to. In the resulting alignment, noncoding portions are characterized by many indels, intercepted by occasional matches. The alignment of noncoding portions of DNA can already be observed in the alignments produced by DIAMOND and thus this side effect does not stem from Patchwork itself. In fact, the Patchwork algorithm will only include noncoding parts if nothing else aligns better to the affected region of the reference sequence.

To mitigate this effect, we have implemented a sliding window-based alignment trimming approach (see figure 2) to rid the alignments from these unwanted regions. This works by scanning the alignment from left to right, cutting all regions where the average distance is below the user-provided distance threshold. The window size and the distance threshold are both set by the user and this entire step can be skipped over in its entirety. This approach tries to avoid cases where a single bad, but correct, match would have otherwise been cut out.

**Fig. 2.**
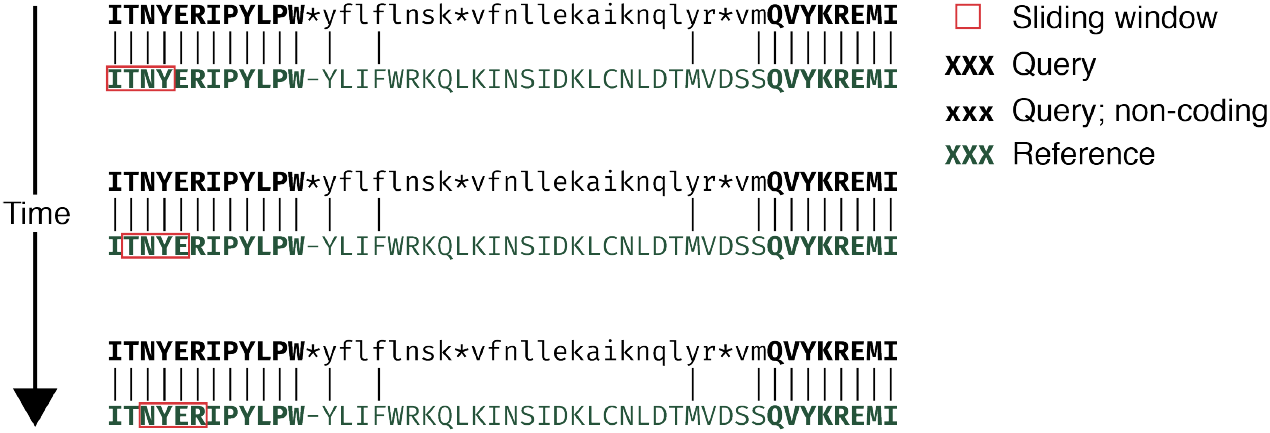
Graphical depiction of the sliding window-based alignment trimming approach. The upper sequence represents the (translated) query sequence aligned to the lower reference sequence, here colored in green. The sliding window, shown in red, moves to the right and removes all residues within, when the average distance falls below the specified threshold. Here, retained residues *after* trimming are shown in a bold font. The putative, noncoding region is shown in lowercase.

### 3.5 Concatenation and realignment of remaining regions

Finally, the resulting set of ordered, non-overlapping sequence regions, are concatenated into one, continuous sequence. The concatenated sequence is then realigned to the reference to obtain the final output sequence and alignment score.

### 3.6 Integrating Patchwork in a phylogenomic pipeline

Most phylogenomic studies include more than a handful taxa and manually concatenating these gets increasingly tedious as the dataset increases in size. Thus, to streamline and simultaneously speed up the downstream analysis, Patchwork includes a set of complementary tools for working with multiple datasets at once. First, `multi_patchwork.sh` can be used to (i) run Patchwork on multiple input files at once and to (ii) concatenate homologous sequences from different taxa into one and the same file(s). The resulting amino acid sequences are in FASTA format and thus the exact downstream analysis used is highly flexible. Nevertheless, a hypothetical bioinformatic pipeline for large-scale phylogenetics may look like the following: (i) BUSCO (Simão et al, 2015; Manni et al, 2021) is used to obtain an initial set of near-universal single-copy orthologs (USCOs) from a (preferably) high-quality genome or transcriptome, sequenced and assembled *de novo* or obtained from a publicly available database; (ii) `multi_patchwork.sh` is used to run Patchwork on a set of assembled genomes, using the USCO-set as a reference, (iii) all resulting markers are aligned using a sequence aligner such as MAFFT (Katoh et al, 2009) or an alignment and alignment trimming program such as GUIDANCE (Penn et al, 2010); finally, (iv) resulting alignments could be filtered for missing data and concatenated into a supermatrix using a tool such as PhyloPyPruner (github.com/fethalen/phylopypruner; Thalén et al. in prep). The resulting data matrix may then be analysed using a multitude of phylogenetic tree inference methods; e.g., maximum likelihood, Bayesian inference, or a coalescent-based methods.

## 4 Benchmarking

To assess the utility and the overall performance of Patchwork, we designed a small study set to recover near-universal single-copy orthologs (USCOs) from genomic Illumina short-read sequence data of the marine annelid *Dimorphilus gyrociliatus,* with the data originally generated as part of a study by Martín-Durán et al (2021). Aside from having an annotated version of the genome publicly available, one of the main advantage of using *D. gyrociliatus* for this study is that the genome is relatively small in size (i.e., 73.82 Mb). As a consequence, assembling a read data for an annelid genome with such a high a high gene density (208.86 genes per Mb) is easier because the read depth and coverage is much higher as a result. However, as we only used short-reads, we created a highly discontinous assembly with low N50 as typical for low-coverage genomic datasets.

This benchmark consists of two phases: In phase 1, we align a shortread-only assembly of *Dimorphilus gyrociliatus* against near-universal single-copy genes found in the long-read, annotated assembly of itself, mentioned above. In phase 2, we instead align the same short-read assembly against USCOs from a chromosome-level assembly of the nereidid *Alitta virens* (GenBank assembly accession: GCA_932294295.1), published and processed by the Wellcome Sanger Institute.

Elapsed time was calculated from the `real` time as reported by the GNU `time` utility, rounded to the nearest second. All analyses were performed on the Ubuntu v20.04 operating system, with a Linux kernel version of 5.13.0, using an Intel^®^ Xeon^®^ Gold 5120 CPU with 28 threads, running at 2.20GHz.

### 4.1 Genome assembly and quality assessment

Raw sequence data was obtained from the Sequence Read Archive (SRA) using the fasterq-dump v2.10.0 tool and the integrity of the data was verified using md5sum v8.30. Quality control (QC) of raw and trimmed reads was performed using FastQC v0.11.9 (https://www.bioinformatics.babraham.ac.uk/) and Trim Galore! v0.6.6 (https://www.bioinformatics.babraham.ac.uk/) was used for automated adapter trimming. The *de novo* assembly was performed using SPAdes v3.15.3 (Nurk et al, 2013), using a K-mer size of 55, and the quality of the assembly was assessed using QUAST v5.0.2 (Gurevich et al, 2013), the results of which are shown in Table 2. We further assessed the quality of the assembly by using BUSCO v5.0.0 (Simão et al, 2015) while utilizing the Metazoa Odb10 dataset, searching a total of 954 BUSCO groups. Out of the 954 BUSCOs, 844 (88.5%) were complete, 822 (86.2%) were complete single-copies, 22 (2.3%) of these were complete and duplicated, 56 (5.9%) were fragmented, and a total number of 54 (5.6%) were missing.

**Table 1.**
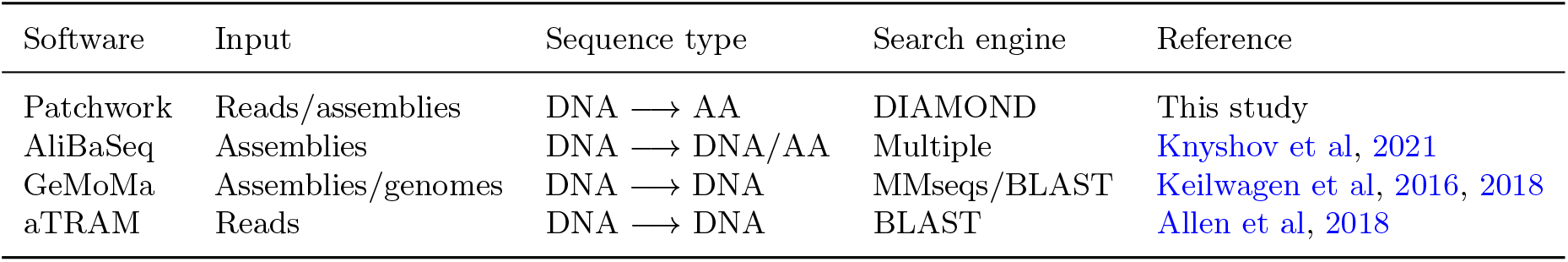
Comparison of software used for evaluation, reproduced from Knyshov et al, 2021.

**Table 2.** Genome assembly quality assessment results of a short-read *de novo* assembly of *Dimorphilus gyrociliatus,* as reported by QUAST (v5.0.2).

| K-mer | No. of contigs | Largest contig | Total length | N50 | L50 |
| --- | --- | --- | --- | --- | --- |
| 55 | 55267 | 682753 | 123971200 | 19851 | 1348 |

### 4.2 *D. gyrociliatus* SPAdes assembly X *D. gyrociliatus* USCOs

We ran our contigs from our *de novo* assembly with sequences from the annelid *Dimorphilus gyrociliatus* through Patchwork v0.5.0, against a pre-annotated protein sequences from the same species (primary accession no. PRJEB37657). The following settings were used for Patchwork: DIAMOND was run using the flag `--iterate` and while discarding any hits with an E-value above 1·10^−3^. Concatenated output sequences that were shorter than 30 AAs were discarded and alignment trimming was performed using a window size of 4 and with a mean minimum required distance of 2.

### 4.3 *D. gyrociliatus* SPAdes assembly X *A. virens* USCOs

Subsequently, we ran the same set of contigs from our *de novo* assembly, made with SPAdes, against a set of near-universal single-copy orthologs (USCOs) from *Alitta virens*. Although both organisms are annelids, they are estimated to have diverged more than 480 million years ago (Dos Reis et al, 2015). In today’s age, one will typically be able to find publicly-available, qualitatively equivalent sequencing data from a more closely-related taxa, but we wanted to see how Patchwork performs on two such distantly-related groups. The chromosome-level assembly of *A. virens* was first obtained from GenBank (assembly accession: GCA_932294295.1). We ran BUSCO v.5.3.1 (Simão et al, 2015), utilizing the “Metazoa” lineage, to recover a total of 897 USCOs from the chromosome-level assembly of *Alitta virens*. For running Patchwork itself, we ran DIAMOND using the flag `--iterate` and while discarding any hits with an E-value above 1·10^−3^. Alignment trimming was performed using a window size of 5 and with a mean minimum distance of −11. No short-sequence filtering was performed in this instance. We subsequently used the `BenchmarkUscos.jl` module from Patchwork to realign the markers we obtained against the pre-annotated set of USCOs in *D. gyrociliatus* itself.

## 5 Results

### 5.1 *D. gyrociliatus* SPAdes assembly X *D. gyrociliatus* USCOs

From the initial 815 markers, we retrieved a set of 788 markers, which corresponds to 96.7% of the total. 27, or 3.3% of the recovered markers were discarded because they were shorter than the 30 AA threshold and 20, or 2.5% of the recovered markers were below 90% in gap-excluded similarity. A detailed summary of these results is shown in table 3. Visualizations of percent identity and query coverage from this run are shown in figure 3. Running Patchwork took a total of 7 minutes and 7 seconds, from start to finish.

**Fig. 3.**
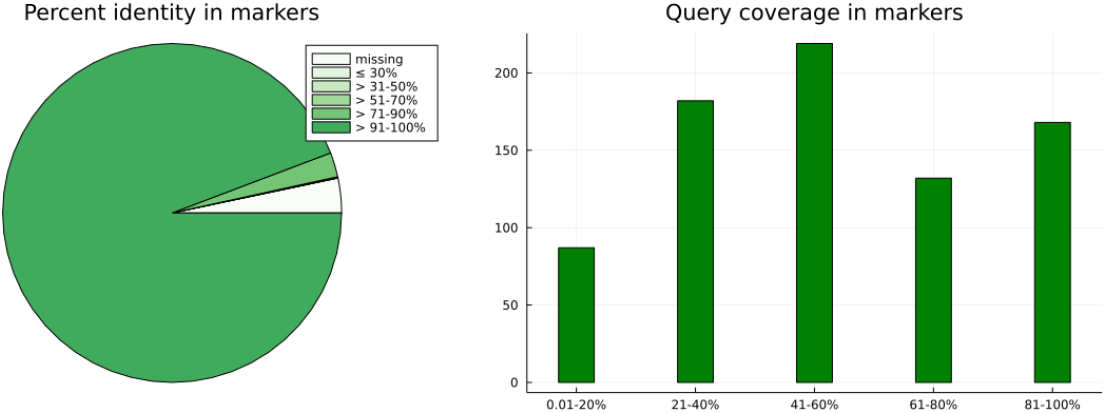
Percent identity-and query coverage in markers based on a Patchwork analysis of a SPAdes assembly of *Dimorphilus gyrociliatus,* targetting 815 single-copy orthologs from itself.

**Table 3.**
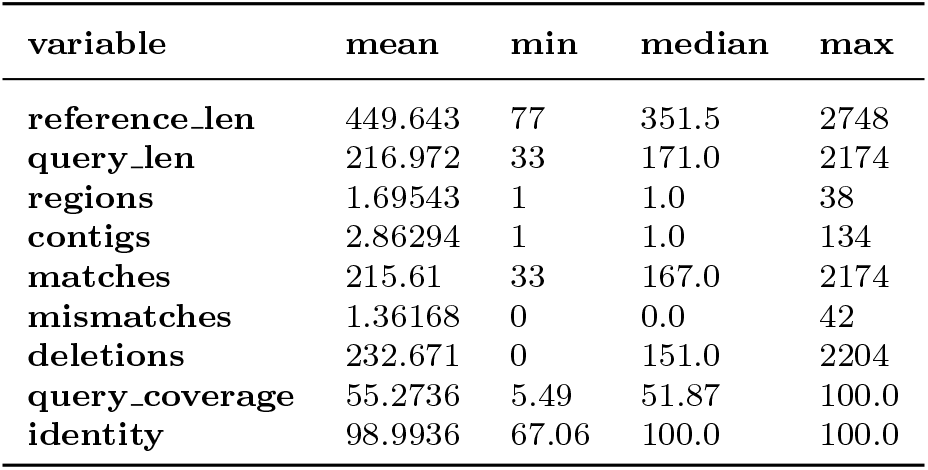
Results from Patchwork when using a *Dimorphilus gyrociliatus* SPAdes assembly as the query and USCOs from a long-read assembly of *Dimorphilus gyrociliatus* as a reference.

| variable | mean | min | median | max |
| --- | --- | --- | --- | --- |
| reference_len | 449.643 | 77 | 351.5 | 2748 |
| query_len | 216.972 | 33 | 171.0 | 2174 |
| regions | 1.69543 | 1 | 1.0 | 38 |
| contigs | 2.86294 | 1 | 1.0 | 134 |
| matches | 215.61 | 33 | 167.0 | 2174 |
| mismatches | 1.36168 | 0 | 0.0 | 42 |
| deletions | 232.671 | 0 | 151.0 | 2204 |
| query_coverage | 55.2736 | 5.49 | 51.87 | 100.0 |
| identity | 98.9936 | 67.06 | 100.0 | 100.0 |

### 5.2 *D. gyrociliatus* SPAdes assembly X *A. virens* USCOs

Out of the 897 *A. virens* USCOs used as a reference, a total of 826 (92.1%) corresponding markers from *D. gyrociliatus* were obtained. 716 of these USCOs successfully aligned back to the pre-annotated set of USCOs found in *D. gyrociliatus.* A detailed summary of these results is shown in table 4. Visualizations of percent identity and query coverage from this run are shown in figure 4. Running *D. gyrociliatus* against the *A. virens* USCOs took a total of 6 minutes and 10 seconds.

**Fig. 4.**
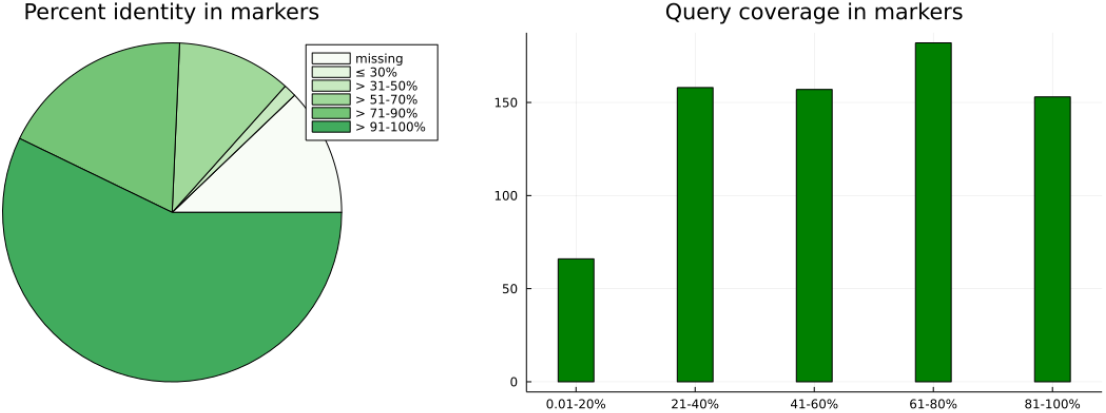
Percent identity-and query coverage in markers based on a Patchwork analysis of a SPAdes assembly of *Dimorphilus gyrociliatus,* targetting a set of 897 near-universal single-copy orthologs (USCOs) from the distantly-related *Alitta virens.* Results shown here are when realigning the recovered USCOs against a set of 826 single-copy orthologs from *D. gyrociliatus* itself.

**Table 4.** Results from Patchwork, using *Dimorphilus gyrociliatus* SPAdes assembly as a query and *Alitta virens* USCOs as a reference, and then re-aligning the resulting query sequences against USCOs from a long-read assembly of *Dimorphilus gyrociliatus.*

| variable | mean | min | median | max |
| --- | --- | --- | --- | --- |
| reference_len | 448.65 | 77 | 358.0 | 2748 |
| query_len | 232.62 | 25 | 172.0 | 2169 |
| no_matches | 197.82 | 25 | 150.5 | 2169 |
| no_mismatches | 28.90 | 0 | 2.0 | 601 |
| no_insertions | 2.15 | 0 | 0.0 | 162 |
| no_deletions | 15.63 | 0 | 4.0 | 355 |
| distance | 970.73 | 127 | 745.0 | 11612 |
| query_cover | 56.88 | 3.29 | 58.925 | 100.0 |
| percent_identity | 89.92 | 33.59 | 97.905 | 100.0 |

## 6 Conclusions

Patchwork is a new software for quickly mining phylogenetic markers from WGS data. Since Patchwork can retrieve homologous regions even in distantly related taxa, this program lends itself especially well for recovering phylogenetic markers for phylogenomic studies, where high-quality transcriptomes are not available for all species of interest. It is simultaneously an efficient way for increasing marker occupancy in poorly assembled genomes and or in the presence of multi-locus exons. Finally, Patchwork allows the user to combine two different data types—transcriptomic and genomic data—into a single dataset, thus further enabling an even larger taxon sampling and encouraging data reusability.

Special consideration should be taken to avoid the creation of chimeric sequences. One way in which such sequences may arise is when orthologous (i.e., genes related via a speciation event) and paralogous (i.e., genes related via a gene duplication event) sequences are merged together. To circumvent this issue, we recommend that the user limit the use of reference sequences to near-universal single-copy orthologs (USCOs). Many programs—e.g., the forementioned program BUSCO—exists for retrieving such sequences from an already assembled genome and these could be used as reference sequences.

The accuracy and the robustness of the results depends on how closely related the two species under study are. The difficulty stems from the ability to accurately predict non-coding regions in aligned contigs; because alignment-trimming relies on gap-excluded identity, choosing the correct cutoff threshold gets increasingly easier as the level of identity approaches 100% (the identity of non-coding regions is likely to stay the same, while the the identity to coding-regions increases). On the upside, high-quality genomes for practically all major lineages exists and are readily available online.

## Data availability

The supplementary data are available at github.com/Animal-Evolution-and-Biodiversity/benchmarking-patchwork.

## Code availability

The source code of Patchwork is available at GitHub (https://github.com/fethalen/patchwork) under the GPLv3 license.

## Acknowledgments

This work was supported by the German Research Foundation (DFG) BL787/8-1 “Phylogenomic analysis of Nereididae (Annelida)”.

## Author’s contributions

F.T. and C.B. conceived of the original design of the software and the study itself. C.K. and F.T. made the software implementation together. F.T. took the lead in writing the manuscript with contributions from C.B. All authors have read and approved of its final version.

## References

Allen JM, Boyd B, Nguyen Np, et al (2017) Phylogenomics from Whole Genome Sequences Using aTRAM. Systematic Biology p syw105. https://doi.org/10.1093/sysbio/syw105

Allen JM, LaFrance R, Folk RA, et al (2018) aTRAM 2.0: An Improved, Flexible Locus Assembler for NGS Data. Evolutionary Bioinformatics Online 14:1176934318774, 546. https://doi.org/10.1177/1176934318774546

Andermann T, Torres Jiménez MF, Matos-Maraví P, et al (2020) A guide to carrying out a phylogenomic target sequence capture project. Frontiers in Genetics 10:1407. https://doi.org/10.3389/fgene.2019.01407, URL https://www.frontiersin.org/article/10.3389/fgene.2019.01407

Andrade SC, Novo M, Kawauchi GY, et al (2015) Articulating “Archiannelids”: Phylogenomics and Annelid Relationships, with Emphasis on Meiofaunal Taxa. Molecular Biology and Evolution 32(11):2860–2875. https://doi.org/10.1093/molbev/msv157

Bezanson J, Edelman A, Karpinski S, et al (2017) Julia: A Fresh Approach to Numerical Computing. SIAM Review 59(1):65–98. https://doi.org/10.1137/141000671

Bragg JG, Potter S, Bi K, et al (2016) Exon capture phylogenomics: Efficacy across scales of divergence. Molecular Ecology Resources 16(5):1059–1068. https://doi.org/10.1111/1755-0998.12449

Buchfink B, Reuter K, Drost HG (2021) Sensitive protein alignments at tree-of-life scale using DIAMOND. Nature Methods 18(4):366–368. https://doi.org/10.1038/s41592-021-01101-x

Call E, Mayer C, Twort V, et al (2021) Museomics: Phylogenomics of the Moth Family Epicopeiidae (Lepidoptera) Using Target Enrichment. Insect Systematics and Diversity 5(2):6. https://doi.org/10.1093/isd/ixaa021

Cronn R, Knaus BJ, Liston A, et al (2012) Targeted enrichment strategies for nextgeneration plant biology. American Journal of Botany 99(2):291–311. https://doi.org/10.3732/ajb.1100356

Dos Reis M, Thawornwattana Y, Angelis K, et al (2015) Uncertainty in the timing of origin of animals and the limits of precision in molecular timescales. Current biology 25(22):2939–2950

Gurevich A, Saveliev V, Vyahhi N, et al (2013) Quast: quality assessment tool for genome assemblies. Bioinformatics 29(8):1072–1075

Heath TA, Hedtke SM, Hillis DM (2008) Taxon sampling and the accuracy of phylogenetic analyses. Journal of Systematics and Evolution p 19

Henikoff JG, Henikoff S (1996) [6] Blocks database and its applications. In: Methods in Enzymology, vol 266. Elsevier, p 88–105, https://doi.org/10.1016/S0076-6879(96)66008-X

Jin JJ, Yu WB, Yang JB, et al (2020) Getorganelle: a fast and versatile toolkit for accurate de novo assembly of organelle genomes. Genome biology 21(1):1–31

Katoh K, Asimenos G, Toh H (2009) Multiple alignment of dna sequences with mafft. In: Bioinformatics for DNA sequence analysis. Springer, p 39–64

Keilwagen J, Wenk M, Erickson JL, et al (2016) Using intron position conservation for homology-based gene prediction. Nucleic Acids Research 44(9):e89–e89. https://doi.org/10.1093/nar/gkw092

Keilwagen J, Hartung F, Paulini M, et al (2018) Combining RNA-seq data and homology-based gene prediction for plants, animals and fungi. BMC Bioinformatics 19(1):189. https://doi.org/10.1186/s12859-018-2203-5

Knyshov A, Gordon ER, Weirauch C (2021) New alignment-based sequence extraction software (ALiBaSeq) and its utility for deep level phylogenetics. PeerJ 9:e11,019. https://doi.org/10.7717/peerj.11019

Lemmon AR, Emme SA, Lemmon EM (2012) Anchored Hybrid Enrichment for Massively High-Throughput Phylogenomics. Systematic Biology 61(5):727–744. https://doi.org/10.1093/sysbio/sys049

Manni M, Berkeley MR, Seppey M, et al (2021) Busco update: novel and streamlined workflows along with broader and deeper phylogenetic coverage for scoring of eukaryotic, prokaryotic, and viral genomes. arXiv preprint arXiv:210611799

Martín-Durán JM, Vellutini BC, Marlétaz F, et al (2021) Conservative route to genome compaction in a miniature annelid. Nature ecology & evolution 5(2):231–242

McCormack JE, Hird SM, Zellmer AJ, et al (2013) Applications of nextgeneration sequencing to phylogeography and phylogenetics. Molecular Phylogenetics and Evolution 66(2):526–538. https://doi.org/https://doi.org/10.1016/j.ympev.2011.12.007, URL https://www.sciencedirect.com/science/article/pii/S1055790311005203, morris Goodman Memorial Symposium

Nurk S, Bankevich A, Antipov D, et al (2013) Assembling genomes and mini-metagenomes from highly chimeric reads. In: Annual International Conference on Research in Computational Molecular Biology, Springer, pp 158–170

Penn O, Privman E, Ashkenazy H, et al (2010) Guidance: a web server for assessing alignment confidence scores. Nucleic acids research 38(suppl_2):W23–W28

Philippe H, Brinkmann H, Copley RR, et al (2011) Acoelomorph flatworms are deuterostomes related to Xenoturbella. Nature 470(7333):255–258. https://doi.org/10.1038/nature09676

Philippe H, de Vienne DM, Ranwez V, et al (2017) Pitfalls in supermatrix phylogenomics. European Journal of Taxonomy (283)

Rhie A, McCarthy SA, Fedrigo O, et al (2021) Towards complete and error-free genome assemblies of all vertebrate species. Nature 592(7856):737–746

Richter S, Schwarz F, Hering L, et al (2015) The utility of genome skimming for phylogenomic analyses as demonstrated for glycerid relationships (annelida, glyceridae). Genome Biology and Evolution 7(12):3443–3462

Rogozin IB, Sverdlov AV, Babenko VN, et al (2005) Analysis of evolution of exon-intron structure of eukaryotic genes. Briefings in Bioinformatics 6(2):118–134. https://doi.org/10.1093/bib/6.2.118, URL https://doi.org/10.1093/bib/6.2.118, https://arxiv.org/abs/https://academic.oup.com/bib/article-pdf/6/2/118/691652/118.pdf

Salzberg SL (2019) Next-generation genome annotation: We still struggle to get it right. Genome Biology 20(1):92. https://doi.org/10.1186/s13059-019-1715-2

Salzberg SL, Phillippy AM, Zimin A, et al (2012) Gage: A critical evaluation of genome assemblies and assembly algorithms. Genome Research 22(3):557–567. https://doi.org/10.1101/gr.131383.111, URL http://genome.cshlp.org/content/22/3/557.abstract, https://arxiv.org/abs/http://genome.cshlp.org/content/22/3/557.full.pdf+html

Sann M, Niehuis O, Peters RS, et al (2018) Phylogenomic analysis of Apoidea sheds new light on the sister group of bees. BMC Evolutionary Biology 18(1):71. https://doi.org/10.1186/s12862-018-1155-8

Schwarz JM, Lüpken R, Seelow D, et al (2021) Novel sequencing technologies and bioinformatic tools for deciphering the non-coding genome. Medizinische Genetik 33(2):133–145. https://doi.org/ doi:10.1515/medgen-2021-2072 URL https://doi.org/10.1515/medgen-2021-2072

Simão FA, Waterhouse RM, Ioannidis P, et al (2015) Busco: assessing genome assembly and annotation completeness with single-copy orthologs. Bioinformatics 31(19):3210–3212

Weigert A, Helm C, Meyer M, et al (2014) Illuminating the Base of the Annelid Tree Using Transcriptomics. Molecular Biology and Evolution 31(6):1391–1401. https://doi.org/10.1093/molbev/msu080

Zhang F, Ding Y, Zhu CD, et al (2019) Phylogenomics from low-coverage wholegenome sequencing. Methods in Ecology and Evolution 10(4):507–517. https://doi.org/10.1111/2041-210X.13145

